# A pool of postnatally-generated interneurons persists in an immature stage in the olfactory bulb

**DOI:** 10.1101/299511

**Authors:** Nuria Benito, Elodie Gaborieau, Alvaro Sanz Diez, Seher Kosar, Louis Foucault, Olivier Raineteau, Didier De Saint Jan

**Author notes:** Corresponding authors Institut des Neurosciences Cellulaires et Intégratives, CNRS UPR 3212, 5 rue Blaise Pascal, 67084 Strasbourg, France, Univ Lyon, Université Claude Bernard Lyon 1, Inserm, Stem Cell and Brain Research Institute U1208, 69500 Bron, France.

## Abstract

Calretinin (CR)-expressing periglomerular (PG) cells are the most abundant interneurons in the glomerular layer of the olfactory bulb. They are predominately generated postnatally from the septal and dorsal sub-ventricular zones that continue producing them well into adulthood. Yet, little is known about their properties and functions. Using transgenic approaches and patch-clamp recording we show that CR(+) PG cells of both septal and dorsal origin have homogenous morphological and electrophysiological properties. They express a surprisingly poor repertoire of voltage-activated channels and fire, at most, one action potential. They also receive little synaptic inputs and NMDA receptors predominate at excitatory synapses. These properties, that resemble those of immature neurons, persist over time and limit the contribution of CR(+) PG cells in network activity. Thus, postnatally-generated CR(+) PG cells continuously supply a pool of latent neurons that unlikely participate in olfactory bulb computation but may provide a so far unsuspected reservoir of plasticity.

## INTRODUCTION

GABAergic interneurons constitute the vast majority of all neurons in the olfactory bulb, the first relay for odor processing in the brain. As elsewhere in the brain, olfactory bulb interneurons are diverse and the functional implications of this diversity are still partially understood (Burton, 2017). Answering this question requires a systematic analysis of each interneuron subtype, covering their developmental origin, morphology, physiology and molecular identity. Many studies have focused on granule cells, the most abundant interneurons in the bulb, that make reciprocal dendrodendritic synapses on mitral and tufted cells lateral dendrites. Much less is known about periglomerular (PG) interneurons that exclusively interact with the apical dendrites of mitral and tufted cells.

PG cells surround each glomerulus where mitral and tufted cells receive synaptic inputs from olfactory sensory neurons (OSN). They are molecularly, morphologically and functionally heterogeneous (McQuiston and Katz, 2001; Hayar et al., 2004; Kosaka and Kosaka, 2007; Panzanelli et al., 2007; Parrish-Aungst et al., 2007; Whitman and Greer, 2007; Najac et al., 2015) and the various subtypes have specific spatial and temporal developmental origins (Batista-Brito et al., 2008; Li et al., 2011; Weinandy et al., 2011). All known PG cell subtypes continue to be generated during the postnatal period, including adulthood, from specific neural stem cells with different embryonic origins and located in defined microdomains in the walls of the subventricular zone (SVZ) (Fiorelli et al., 2015). CR(+) PG cells are the most abundant PG cell subtype, representing 40-50% of the whole PG cell population. They are 2-3 times more numerous than other subtypes expressing calbindin (CB), parvalbumin or tyrosine hydroxylase (TH) (Kosaka and Kosaka, 2007; Panzanelli et al., 2007; Parrish-Aungst et al., 2007; Whitman and Greer, 2007). CR(+) PG cells are also the most abundant PG cells generated postnatally. Contrasting with CB(+) and TH(+) cells whose production is maximal during embryogenesis and declines after birth, production of CR(+) PG cells peaks around birth and continues throughout life (De Marchis et al., 2007; Ninkovic et al., 2007; Batista-Brito et al., 2008; Li et al., 2011; Weinandy et al., 2011).

Functionally, PG cells have been classified into two classes: type 1 PG cells that receive direct synaptic inputs from OSNs, and type 2 PG cells that do not. Like CB(+) PG cells, dendrites of CR(+) PG cells avoid compartments of the glomerulus that contain OSN terminals and have therefore been classified as type 2 PG cells (Kosaka and Kosaka, 2007). Consistent with this classification, we recently confirmed that CR(+) PG cells do not receive synaptic inputs from OSNs (Najac et al., 2015). Several observations, however, suggest unique properties of CR(+) PG cells. For instance, they have different embryonic origins than CB(+) PG cells (Kohwi et al., 2007) and their development is regulated by specific transcription factors (Waclaw et al., 2006; Kohwi et al., 2007; Qin et al., 2017; Tiveron et al., 2017). They also present distinctive electrical membrane properties (Fogli Iseppe et al., 2016).

We combined transgenic approaches and patch-clamp recording in olfactory bulb slices to examine the origin, intrinsic and synaptic properties of this so far overlooked interneuron subtype. We found that CR(+) PG cells show a dual spatial origin, i.e. the septal and dorsal SVZ that continue producing them well into adulthood, although at different pace. In spite of these distinct origins, CR(+) PG cells are homogeneous in term of morphological and electrophysiological properties. Interestingly, CR(+) PG cells show immature traits which persist over extended periods of time and limit their contribution to olfactory information processing. Our data instead suggest that CR(+) PG cells constitute a pool of latent neurons that may provide a previously unsuspected reservoir of plasticity.

## RESULTS

### 1 CR(+) PG cells from different spatial origin are morphologically homogeneous

The postnatal SVZ produces distinct subtypes of PG interneurons in a region dependent manner. We have shown that CR(+) PG cells originate from the dorsal and septal most regions of the SVZ at birth (Fernandez et al., 2011). The temporal dynamic of their postnatal production, however, remains elusive. To address this question, we combined targeted EPO of specific SVZ microdomains with a Cre-lox approach in P2-old Rosa^YFP^ mice for permanent labeling and fate mapping of neural stem cells located in the lateral, dorsal and septal aspects of the postnatal SVZ (**Fig 1A**). We sacrificed the mice at 1.5 months and 3 months and quantified recombined CR(+) PG cells expressing the immature neuronal marker doublecortin (DCX) to determine the spatial origin of cells generated in the last 2/3 weeks (**Fig 1B**). CR(+) PG cells constituted the majority of the DCX(+) PG cells derived from the septal and, to a lesser extent, dorsal SVZ (83.1±6.2% and 59.5±10.3%, respectively, at 1.5 months and 88.3±2.9% and 44.7±17.6%, respectively, at 3 months; >50 YFP(+)/DCX(+) cells per EPO in n=3 mice at both time points), whereas the lateral SVZ rarely produced CR(+) PG cells. Thus, both the septal and dorsal microdomains remain active in producing CR(+) PG interneurons in adulthood, in similar proportions as previously observed at birth (Fernandez et al., 2011).

**Figure 1.**
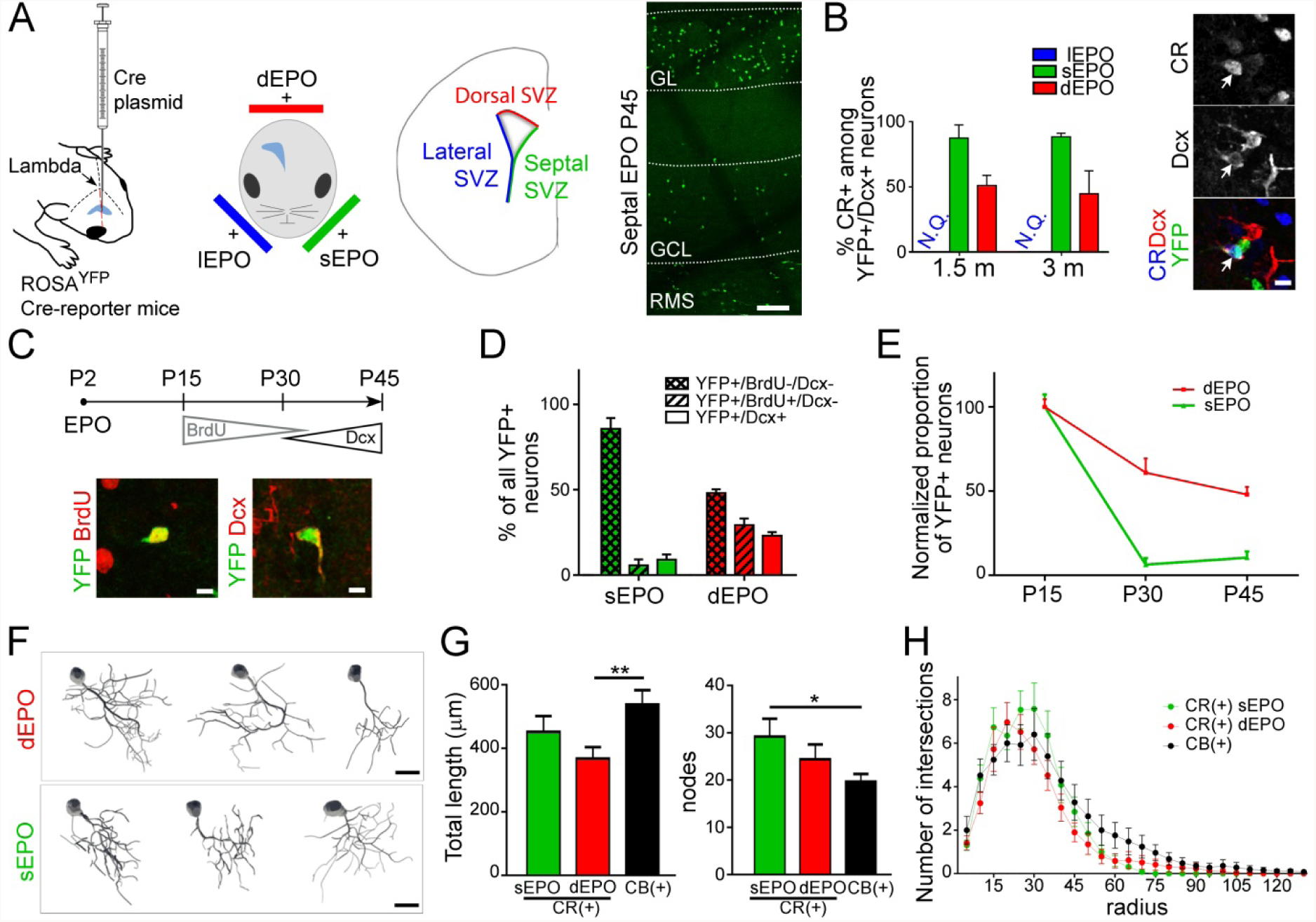
Spatio-temporal origin and morphology of postnatally generated CR(+) PG interneurons. **A)** We modified a previously established targeted EPO approach for permanent labeling and fate mapping of neural stem cells that reside in defined microdomains of the postnatal SVZ. A plasmid expressing the Cre recombinase was electroporated in dorsal, lateral and septal SVZ of Rosa^YFP^ Cre-reporter mice allowing the recombination and permanent expression of a yellow fluorescent protein (YFP) in neural stem cells and their progeny. The picture on the right shows the accumulation of YFP(+) cells in the olfactory bulb 45 days after a septal EPO. Scale bar: 100 µm. **B)** Quantification of the fraction of CR(+) PG interneurons among DCX(+)/YFP(+) PG cells at 1.5 and 3 months reveal their stable spatial origin over time. The number of cells expressing CR was negligible following lateral electroporation, and was not quantified. Scale bar: 5 µm. **C)** Long term fate mapping was combined with immunodetection of BrdU and DCX to discriminate between cohorts of neurons sequentially produced at distinct postnatal times over a period of 1.5 months. Insert shows representative images of YFP(+) cells expressing BrdU or DCX. Scale bars: 5 µm. **D-E)** Quantification of YFP(+) PG cells born at early (BrdU-/DCX-), intermediate (BrdU+/DCX-) and late time (BrdU-/ DCX+) during the first 1.5 months of postnatal life reveal the temporal dynamic of neurogenesis from the septal and dorsal SVZ microdomains. Data were normalized in E for better clarity. **F)** Neurolucida reconstruction of YFP(+)/CR(+) PG interneurons originating from the dorsal and septal SVZ at 21 days post-EPO. **G)** Quantification of dendritic arborisation length and number of nodes in CR(+) PG interneurons originating from the dorsal (red) and septal (green) SVZ. Values for CB(+) PG cells reconstructed at 21 days post dorsal EPO are given for comparison. Scale bars: 10 µm. **H)** Sholl analysis representing the number of dendrite intersections with concentric circles of gradually increasing radius. Abbreviations: EPO: electroporation; Cre: Cre recombinase; SVZ: subventricular zone; CR: Calretinin; CB: calbindin; PG: periglomerular; GL: glomerular layer; GCL: granule cell layer; RMS: rostral migratory stream; N.Q.: not quantified.

We next assessed the temporal dynamic in neurogenic activity of the septal and dorsal SVZ microdomains. We administrated BrdU through drinking water during the second half of the first month of postnatal life, and sacrificed the animals at 1.5 months (**Fig 1C**). BrdU and DCX immunodetection allowed us to discriminate early (BrdU(-)/DCX(-)), intermediate (BrdU(+)/DCX(-)) and late born interneurons (BrdU(-)/DCX(+)) among recombined YFP(+) cells (**Fig 1C**) (n=164 YFP(+) cells in 6 mice for dorsal EPO; n=240 YFP(+) cells in 3 mice for septal EPO). This analysis revealed a gradual decrease of neurogenesis for both microdomains, which appeared more pronounced for the septal SVZ (**Fig 1D, E**). Thus, while CR(+) PG interneurons present a dual origin, the contribution of the septal SVZ declines faster than the contribution of the dorsal SVZ in their generation.

These observations prompted us to investigate possible morphological differences between CR(+) PG cells produced by the septal and dorsal SVZ. Individual CR(+) PG cells originating from both SVZ microdomains were reconstructed 21 days post-EPO (**Fig 1F**) and compared with CB(+) PG cells at the same age. All reconstructed cells (n=26 CR(+) from septal EPO, n=29 CR(+) and n=25 CB(+) from dorsal EPO, in n=3 mice per condition) had the typical morphology of mature PG cells i.e. no axon and a polarized dendritic tree ramifying within a single glomerulus (**Fig 1F**). Quantitative analysis of dendritic length and number of nodes (**Fig 1G**), as well as a Sholl analysis that measures the number of times the dendrites intersect with consecutive concentric circles (**Fig 1H**) revealed a similarly complex and a rather homogeneous morphology for CR(+) PG cells, independent of their origin (p>0.05 for all dorsal/septal CR(+) comparison in **Fig 1G** and **H**). The Sholl analysis also revealed no significant difference between CR and CB-expressing PG cells (**Fig 1H**), although dorsal CR(+) cells had a tendency for shorter dendrites and septal CR(+) cell a tendency for more nodes **(Fig 1G)**. Thus, CR(+) PG cells with distinct spatial origin have largely similar morphologies and, despite small differences that may reflect the morphological immaturity of some cells included in this analysis, resemble CB(+) PG cells.

### 2 CR::EGFP transgenic mice recapitulate the complex spatio-temporal origin of CR(+) PG cells

The CR::EGFP transgenic mouse expresses EGFP under the control of the CR promoter (Caputi et al., 2009). In the glomerular layer of the olfactory bulb, EGFP is exclusively expressed in CR(+) PG cells (Najac et al., 2015)(**Fig 2A**). To assess if EGFP labels a subpopulation of CR(+) PG cells with a specific origin, we transiently electroporated the septal or dorsal SVZ of CR::EGFP newborn pups with a plasmid encoding tdTomato (Tom) **(Fig 2A)**. Three weeks later, we quantified the fraction of Tom(+)/CR(+) PG cells that also expressed EGFP (n=69 and n=43 Tom(+)/CR(+) PG cells counted in 3 mice, respectively). We found that both the septal and dorsal SVZ produced EGFP(+) PG cells, although in different proportion (67±10% and 29±4%, respectively; **Fig 2B**). The number of EGFP(+) among CR-expressing PG cells slightly decreased from ~80% at 10 days to ~50% in adult mice (**Fig 2C**), possibly reflecting the increasing contribution of the dorsal SVZ to their production. However, reconstructions of individual Tom(+)/CR(+) cells positive (n=43) or negative (n=12) for EGFP revealed no morphological differences between the two populations (p=0.58 for total length and p=0.91 for surface; **Fig 2D, E**).

**Figure 2.**
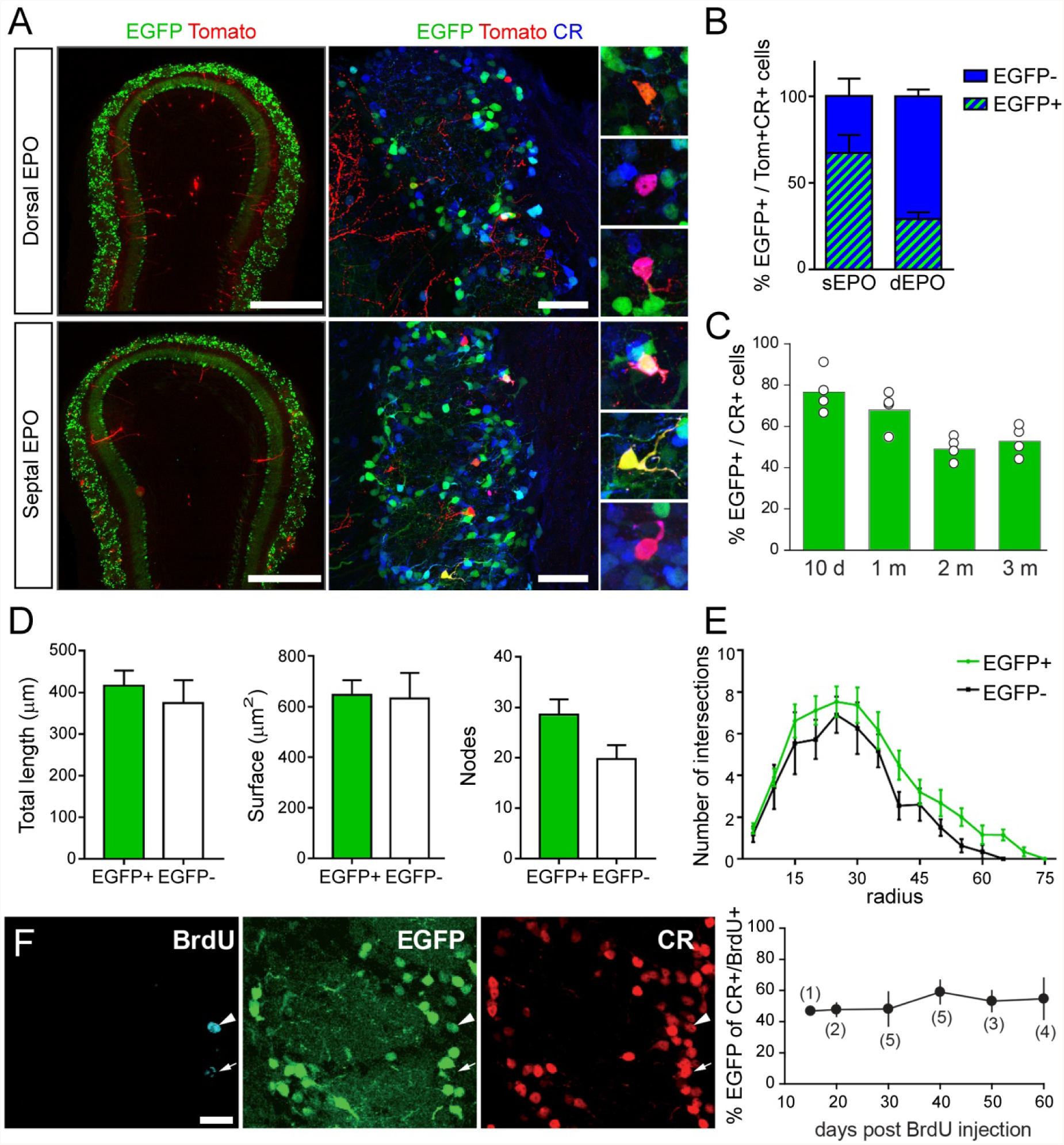
Labeling of a representative population of CR(+) PG interneurons in the CR::EGFP transgenic mice. **A)** Fate mapping of EGFP(+)/CR(+) PG interneurons from dorsal and septal origin by EPO of a tdTomato expressing plasmid in CR::EGFP transgenic mice. Captions show marker expression by select cells: Tom(+)/CR(+) appear in orange/yellow, while those also expressing EGFP appear in pink. Scale bars: 500 µm (left panels) and 50 µm (right panels). **B)** Proportion of CR(+) PG interneurons expressing EGFP following dorsal and septal EPO. Note that EGFP expression can be observed in both populations of CR(+) PG interneurons. **C)** Percentage of EGFP expression among the CR(+) PG population at different postnatal ages. **D)** Neurolucida reconstruction of EGFP(+) and EGFP(-) Tom(+)/CR(+) PG interneurons reveals comparable dendritic arborisation length, surface and number of nodes. **E)** Sholl analysis representing the number and distribution of dendrite intersections with concentric circles of gradually increasing radius. **F)** Birth dating of CR(+) PG interneuron demonstrates stable EGFP expression in CR(+) PG interneurons up to 60 days after their generation. Scale bar: 10 µm

Finally, we conducted a birth dating experiment to determine if the EGFP labeling also recapitulates the age diversity of CR(+) PG cells or only transiently labels CR(+) PG cells at a specific period of maturation. BrdU was administered to 30 days-old CR::EGFP mice (n=20) to label cells generated at that age. Mice were then sacrificed at different intervals (15-60 days) after the injection to evaluate the time window during which BrdU-expressing CR(+) PG cells express EGFP. This analysis revealed that the percentage of BrdU(+)/EGFP(+) PG cells among BrdU(+)/CR(+) PG cells (around 50%) remained stable up to 60 days post-injection, thereby excluding a transient expression of EGFP in CR(+) PG cells. Thus, EGFP expression is not transient and labels a fraction of CR(+) PG cells that can be anywhere between 15 and, at least, 60 days old (**Fig 2F**). Together, these data indicate that EGFP stably labels a population of CR(+) PG cells that includes a representative population of neurons, from different spatial origin and at different stages of maturation.

### 3 CR(+) PG cells fire at most one action potential at any stage of maturation

Having established that EGFP expression faithfully reflects the spatio-temporal diversity of CR(+) PG cells, we examined the functional properties of EGFP(+) PG cells using patch-clamp recording in acute olfactory bulb horizontal slices from CR::EGFP mice. First, we examined the intrinsic membrane properties of EGFP(+) PG cells using whole-cell recordings. EGFP(+) PG cells had an electrical membrane resistance of 2.0 ± 0.12 GOhm (n=74). This high membrane resistance, which is most likely underestimated because of the current leak through the pipette seal, suggests the expression of few ionic channels. In the current-clamp mode, depolarizing current injections from a holding potential maintained around −70 mV induced at most a single action potential, sometime followed by a spikelet (**Fig 3A)**. Further depolarizing the membrane did not elicit additional action potentials and, instead, passively depolarized the cell to positive potentials until the end of the step. This passive depolarization was seen in every cell tested and was therefore a robust functional marker of CR(+) PG cells. Only 60% of the cells (n=47/77) fired an overshooting action potential (with an amplitude >40 mV; **Fig 3B**). The other cells did not fire at all or fired a single rudimentary action potential (amplitude <40 mV, n=30), much like immature newborn granule cells in the olfactory bulb (Carleton et al., 2003) or in the hippocampus (Overstreet et al., 2004; Esposito et al., 2005). Three examples of this diversity are shown in **Fig 3A**. EGFP(+) PG cells firing a full size action potential had more complex voltage responses and significantly faster membrane time constant (14.1 ± 1.4 ms for n=13 cells with spike height >65 mV vs. 26.6 ± 1.8 ms for n=13 cells with spike height <20 mV, p<0.0001), consistent with a maturation of the membrane properties. In the most mature cells, the action potential was sometime riding on the top of a calcium spike, and hyperpolarizing steps evoked a sag in membrane potential suggesting the activation of an Ih current. In rare cases, we recorded cells that spontaneously and regularly fired a single action potential at about 0.5-1 Hz (not shown). This cell to cell variability, that can be illustrated using the size of the spike as an indicator of cell maturation, was often seen within the same slice from animals at different developmental ages (P15-P38; **Fig 3C**). Thus, consistent with their ongoing production, CR(+) PG cells constitute a population of neurons with variable membrane properties that likely reflect different stages of maturation.

**Figure 3:**
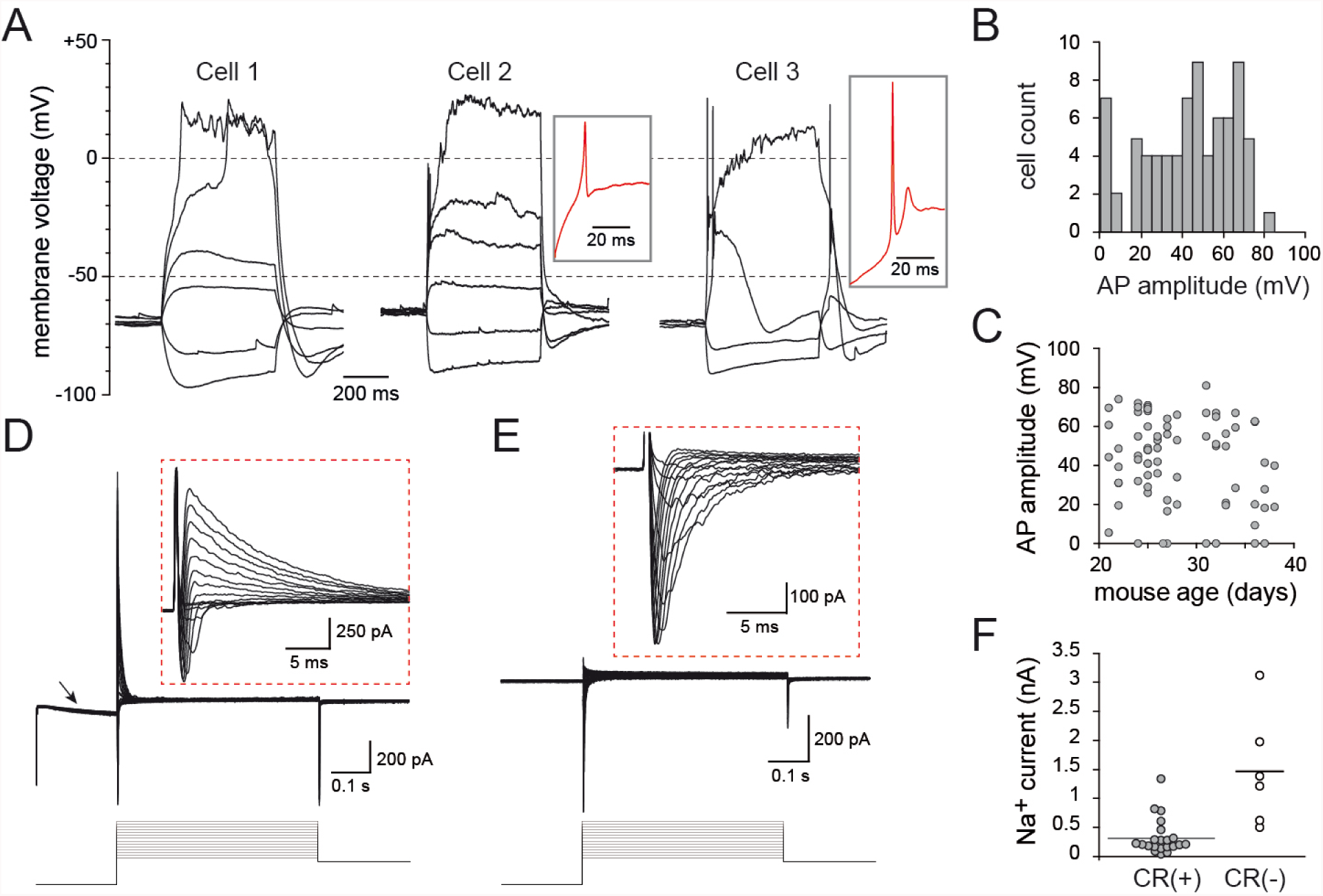
Intrinsic membrane properties of CR(+)/EGFP(+) PG cells. **A)** Voltage responses of three representative CR(+) PG cells to 500 ms long current steps. Cell 1 did not fire, cell 2 fired a small non overshooting action potential and cell 3 fired a single overshooting action potential followed by a spikelet. Red traces in insets are a zoom on the first suprathreshold response of cell 2 and 3. **B)** Distribution histogram of action potential amplitudes across all the EGFP(+) PG cells tested. **C)** The amplitude of the action potential did not depend on the age of the mouse. **D)** Depolarizing voltage steps (5mV/500 ms) induced fast inward Na^+^ currents and outward K^+^ currents (shown at a higher magnification in the inset). The hyperpolarizing prepulse (−40 mV from Vh=-75 mV) evoked an Ih current (arrow). **E)** Na^+^ currents were isolated in the presence of K^+^ and Ca^2+^ channel blockers (Cs, TEA, 4-AP and Cd). **F)** Maximal Na^+^ current amplitudes recorded in CR(+)/EGFP(+) PG cells compared with those recorded in CR(-) PG cells in the same slices. Recordings were done as in (E).

Next, we examined the voltage-activated conductances underlying the membrane potential responses. Consistent with a recently published study (Fogli Iseppe et al., 2016), classical voltage-clamp protocols activated few conductances and, remarkably, no delayed K+ currents. While inward Ih currents were often induced by a 40 mV hyperpolarizing pre-pulse, depolarizing steps only evoked fast inward Na^+^ currents followed by rapidly inactivating outward K^+^ currents (**Fig 3D**). Ca^2+^ inward currents were not seen under these recording conditions suggesting that they may be too small to be detected (see (Fogli Iseppe et al., 2016)). K^+^ currents were large (1 ± 0.1 nA at Vh=-10 mV) and fast (deactivation time constant: 12.4 ± 1.0 ms, n=19) in every cell tested, and strongly reduced by a 20 mV depolarizing prepulse (88 ± 2 % decrease, n=9, not shown), consistent with A-type K^+^ currents. Na^+^ currents were, in contrast, often small or totally absent in some cells (n=21 cells, **Fig 3E-F**) and only a fraction of the cells expressed Na^+^ currents comparable in size to those seen in other non-labeled PG cell subtypes. Together, these data indicate that EGFP(+) PG cells, regardless of their degree of maturation, express a surprisingly poor repertoire of voltage-activated conductances.

### 4 CR(+) PG cells receive little synaptic inputs at immature synapses

We have already shown that EGFP(+) PG cells receive less spontaneous excitatory synaptic inputs (sEPSC) than other type 2 PG cells and respond to a stimulation of the olfactory nerve with a smaller composite EPSC (Najac et al., 2015). This is here illustrated with a paired recording of two PG cells, one EGFP(+) and one EGFP(-), projecting into the same glomerulus (**Fig 4A**). To complement our previous analysis, we recorded sEPSC from animals at different ages (n=52 cells) and also examined their spontaneous inhibitory inputs (sIPSC, n=42 cells). The majority of the cells tested received a low frequency of spontaneous EPSCs (90% received less than 5 sEPSC/s and 75% less than 2 sEPSC/sec; **Fig 4B**). This frequency is much lower than in other type 2 PG cells. For instance, PG cells labeled in the Kv3.1-EYFP reporter mouse on average receive 17 sEPSC/s (Najac et al., 2015). EGFP(+) cells also received few spontaneous IPSC (90% less than 5 sIPSC/s and nearly 70% less than 2 sIPSC/s). There was no obvious relationship between the frequencies of sEPSC and sIPSC in cells in which these two synaptic inputs were recorded (not shown). There was also no obvious relationship between the age of the animal and the synaptic input frequencies in EGFP(+) PG cells (**Fig.4B**). Remarkably, cells receiving no synaptic input were observed at all ages.

**Figure 4:**
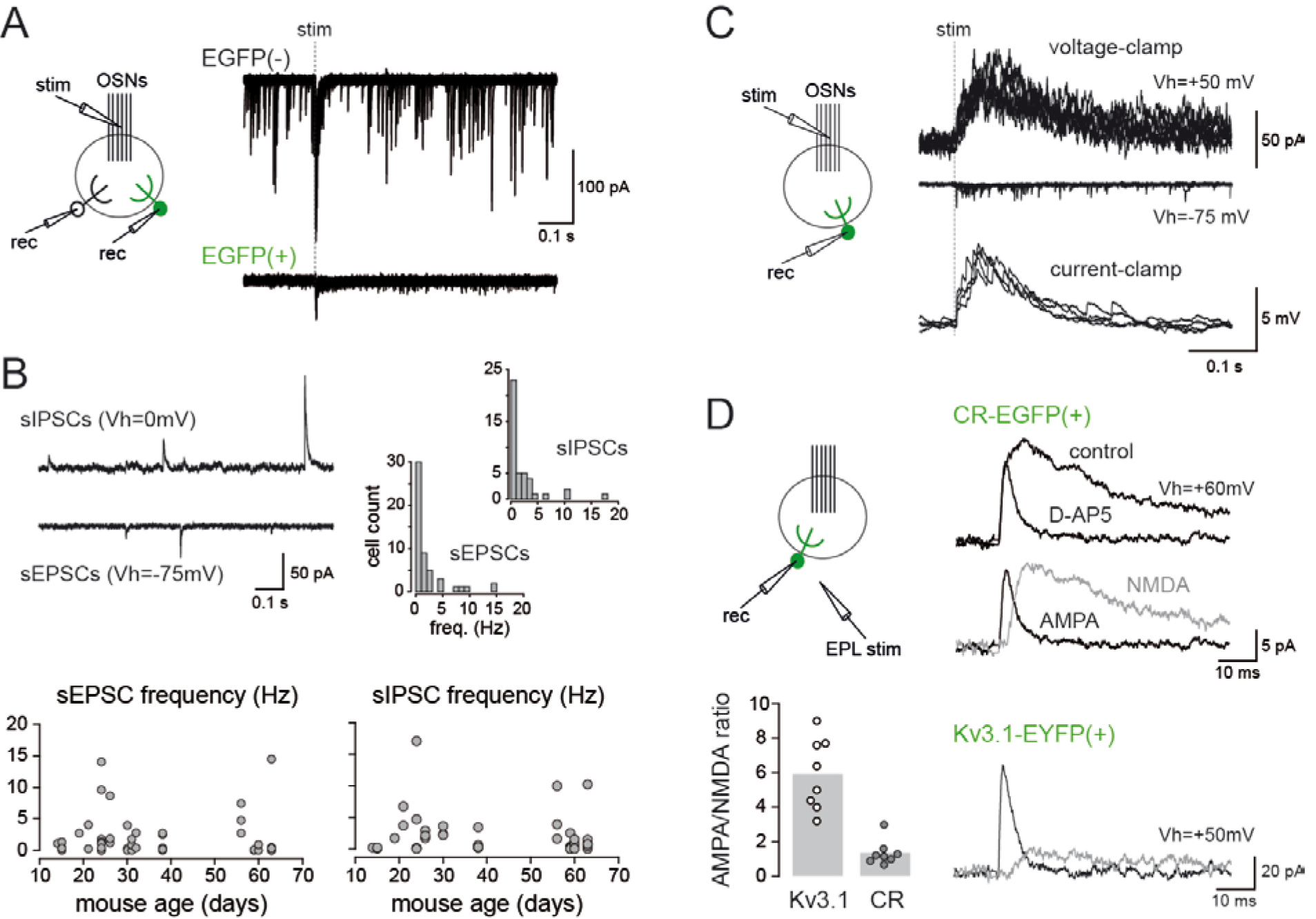
Synaptic inputs of CR(+)/EGFP(+) PG cells. **A)** Paired recording between an EGFP(+)/CR(+) PG cell and a non-labeled PG cell projecting into the same glomerulus. 15 traces are superimposed for each cell. The dashed line indicates OSN stimulation. **B)** Spontaneous IPSCs (sIPSCs recorded at Vh=0 mV, top trace) and EPSCs (sEPSCs recorded at Vh=-75 mV, lower trace) in a representative EGFP(+)/CR(+) PG cell. Right, distribution histograms (bin of 1 Hz) of synaptic input frequencies showing the predominance of cells receiving few inputs. Bottom, summary graphs of sEPSCs and sIPSCs frequencies in cells from mice at different developmental ages. Each dot represents a cell. **C)** OSN-evoked synaptic responses of an EGFP(+)/CR(+) PG cell recorded in the voltage-clamp mode at a positive holding potential (top traces) or at a negative potential (middle traces), and in the current-clamp mode (bottom traces). Several traces are superimposed each time. Stimulation 150 µA. **D)** Comparison of AMPA/NMDA ratios at excitatory synapses in CR-EGFP(+) PG cells (top traces) and in Kv3.1-EYFP(+) PG cells (bottom traces). EPSCs were elicited by an electrical stimulation in the EPL and recorded at a positive holding potential in control conditions and in the presence of D-AP5 (50 µM). The NMDA component (grey trace) was obtained by subtracting the D-AP5 trace from the control trace. Summary bars show the average AMPA/NMDA ratio in CR(+)/EGFP(+) PG cells and in Kv3.1-EYFP(+) PG cells. Each dot represents a cell.

Stimulation of the OSN evoked few fast asynchronized EPSCs in voltage-clamp recordings, consistent with the small number of connections with mitral and tufted cells. Yet, these small EPSCs (average amplitude 12 ± 3 pA at Vh = −75 mV, n=5) converted into slow summating EPSPs and produced a significant depolarization in the current-clamp mode (6 ± 1 mV, n=5; **Fig 4C**). Interestingly, the slow time course of the OSN-evoked EPSP was similar as the time course of the OSN-evoked EPSC recorded in voltage-clamp at a positive holding potential, suggesting a strong contribution of NMDA receptors (**Fig 4C**). To further test this hypothesis, we determined the contribution of NMDA receptors in individual EPSCs evoked by a stimulation of mitral and tufted cells in the external plexiform layer (EPL). Evoked EPSCs were recorded at a positive holding potential (Vh = +50/60 mV), first in control condition and then in the presence of the NMDA receptor antagonist D-AP5 (50 µM) to isolate the AMPA component. Then, we quantified the AMPA/NMDA ratio of the individual EPSC (**Fig 4D**). Under these conditions, the AMPA/NMDA ratio at mitral/tufted cells to CR+ PG cell synapses was 1.4 ± 0.2 (n=8 EGFP(+) PG cells, all firing an overshooting action potential with amplitude range 42-67 mV). This AMPA/NMDA ratio was several folds lower than in PG cells labeled in the Kv3.1-EYFP reporter mouse (AMPA/NMDA ratio: 5.9 ± 0.7, n=8 EYFP(+) PG cells, t(14)=5.9, p<0.0001)(**Fig 4D**). This confirms the high density of NMDA receptors at excitatory synapses on CR(+) PG cells, a feature of immature synapses.

### 5 Synaptic integration of CR(+) PG cells does not progress over time

The diversity of membrane properties observed in our random sampling of EGFP(+) PG cells is consistent with a developmental maturation of the intrinsic membrane properties across cells generated at different time. How synaptic integration progress in time is less clear. To clarify this question, we examined the properties of age-matched EGFP(+) PG cells co-labeled, using targeted EPO of the septal SVZ, with tdTomato (**Fig 5A**). Tom(+)/EGFP(+) PG cells were recorded at different intervals after EPO (n=50 cells, 15-63 days post EPO). As expected, these cells had similar properties as those described before, i.e. they fired at most a single action potential and received little spontaneous synaptic inputs. However, most of the cells tested fired an overshooting action potential (n=22/25 at ≥20 days post EPO; **Fig 5B**) suggesting that cells that did not fire or fired a rudimentary action potential in our random sampling of EGFP(+) PG cells were <20 days old. In contrast, the frequency of the spontaneous synaptic inputs was not correlated with the age of the cell (**Fig 5C**). Similar to randomly chosen EGFP(+) PG cells, Tom(+)/EGFP(+) PG cells received between 0 and 10 EPSCs per second (mean 2.3 ± 2.7, n=47) and, for the majority of them, less than 3 IPSCs/s (mean 1.0 ± 1.7, n=41) whether they were young (15-23 days post EPO) or older (54-63 days post EPO). Thus, the mean frequency of sEPSC was not different in young cells (2.2 ± 0.5 EPSC/sec, n=25) and in old cells (2.4 ± 0.8 EPSC/sec, n=12, p=0.88). The mean sIPSC frequency was also not different in young cells (0.7 ± 0.2 IPSC/sec, n=21) and in old cells (1.2 ± 0.3 IPSC/sec, n=11, p=0.17). Importantly, cells without any detectable synaptic inputs were recorded at all ages. Thus, synaptic integration of CR(+) PG cells does not progress over time.

**Figure 5:**
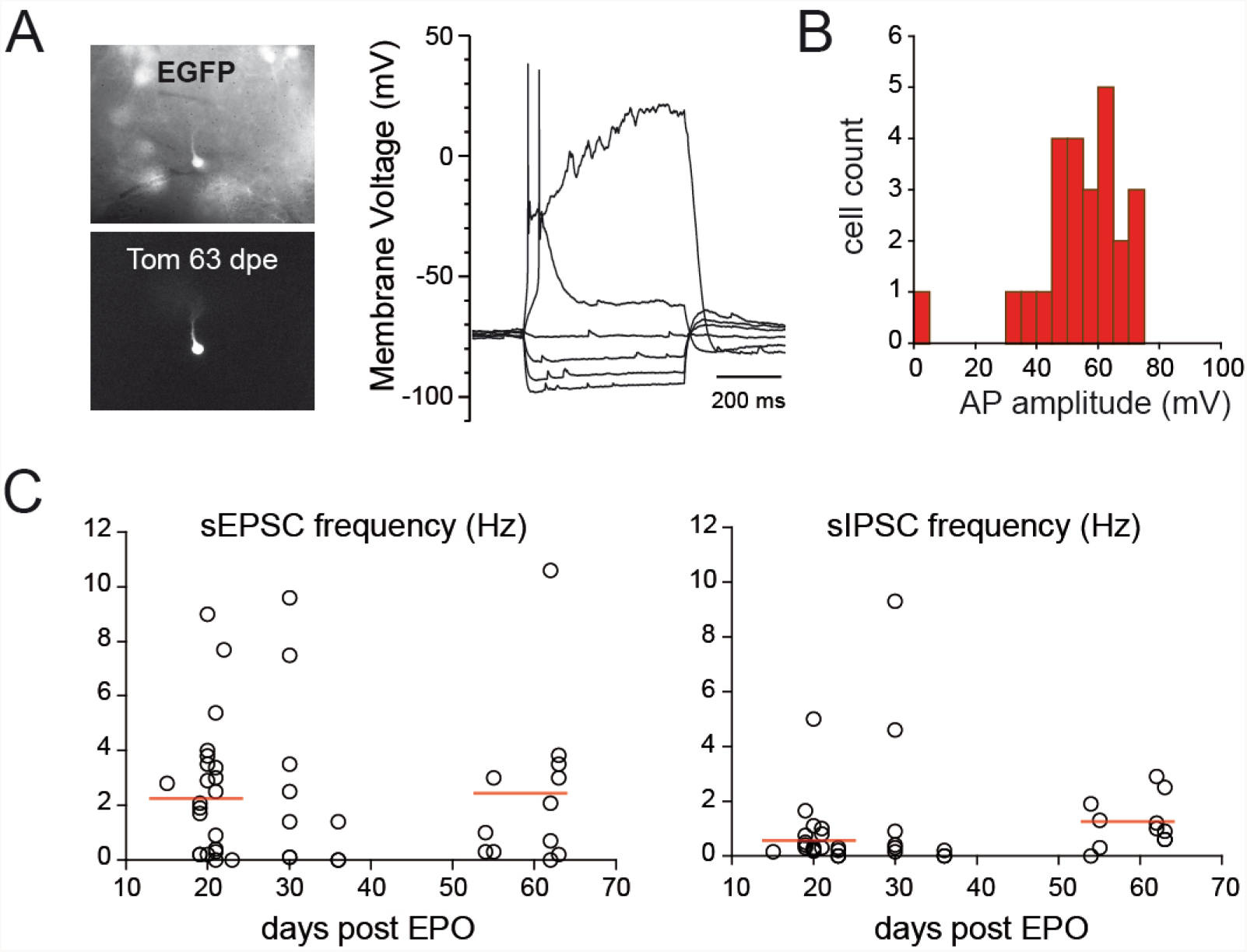
Synaptic integration of CR(+)/EGFP(+) PG cells does not change over time. **A)** Current-voltage relationship of an EGFP(+) PG cell (top) co-labeled with tdTomato (Tom, bottom), 63 days after EPO of a tdTomato-expressing plasmid in a CR::EGFP transgenic mouse. **B)** Distribution of action potential amplitudes in all the EGFP(+)/Tom(+) cells recorded 19-63 days post-EPO. **C)** Excitatory (left) and inhibitory (right) input frequencies in EGFP+/Tom+ PG cells recorded at different intervals after EPO. Each dot represents a cell. Horizontal bars show average frequencies for cells recorded at 15-23 days or 54-63 days post EPO.

### 6 CR(+) PG cells are less recruited than other PG cells

Low innervation counteracts high intrinsic membrane excitability and prevents the firing of newly-generated immature neurons (Dieni et al., 2016). In the other hand, temporal summation of small, input-specific, NMDA receptor-mediated EPSPs converts into reliable spiking in immature neurons (Li et al., 2017). Thus, sparse activation of immature neurons may play important and specific functions in circuit computation. We examined the afferent-driven recruitment of CR(+) PG cells using paired loose cell-attached recording. We compared the OSN-evoked firing activity of EGFP(+) PG cells with those of other PG cell subtypes embedded in the same network. Stimulation intensities were first adjusted in order to reliably elicit the discharge of a randomly chosen EGFP(-) PG cell. Then, we simultaneously recorded from an EGFP(+) cell projecting into the same glomerulus (n=18 pairs in slices from P31-P39 old mice). As illustrated in **Fig.6A**, spontaneous and evoked action potential capacitive currents were classically biphasic and large in EGFP(-) PG cells (average amplitude 121 ± 28 pA, range 18-532 pA) (**Fig 6C**). The number of spikes evoked by the stimulation of the OSNs (from 2 to >16, average 5.8 ± 4), as well as the duration of the burst varied across cells, consistent with the diverse functional profiles of PG cells (**Fig 6A,B**) (Najac et al., 2015). In sharp contrast, OSN stimulations produced no response at all (n=8 cells) or a single, occasionally two, small monophasic capacitive currents (15 ± 2 pA, n=10 cells) in EGFP(+) PG cells (**Fig 6A-C**). Increasing the intensity of stimulation did not change the responses indicating that EGFP(+) PG cells do not have a higher activation threshold. Moreover, spontaneous spikes were rarely seen in EGFP(+) cells (few events seen in only 3/18 cells) whereas they were frequent and seen in the majority of EGFP(-) cells (11/18). Thus, CR(+) PG cells are remarkably silent compared with other PG cell subtypes and, in sharp contrast with CB(+) PG cells (Najac et al., 2015), are poorly recruited by afferent inputs.

**Figure 6:**
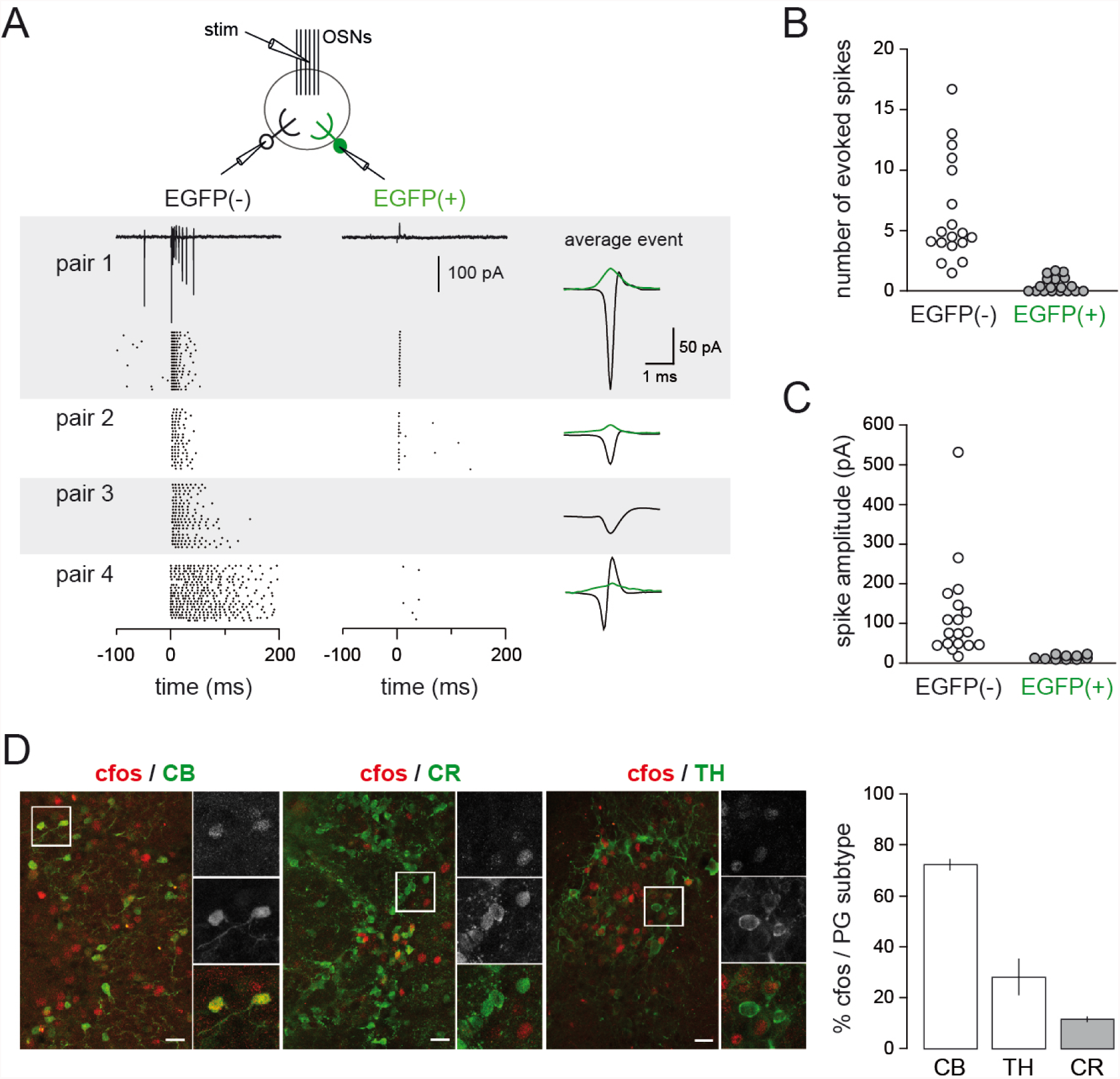
CR(+) PG cells are poorly recruited *in vitro* and *in vivo*. **A)** OSN-driven spiking of EGFP(+)/CR(+) PG cells in olfactory bulb slices. Two PG cells projecting into the same glomerulus, one EGFP(+) and one EGFP(-), were recorded in the loose-cell attached configuration. Their firing was elicited by an electrical stimulation of the OSNs. Raster plots from 4 pairs are shown with 20 consecutive sweeps for each pair. The EGFP(-) cell is on the left, the EGFP(+) on the right. A typical response is shown for pair 1. OSNs were stimulated at t=0, at an intensity of 30-200 µA (stimulation artifact has been blanked). The average evoked capacitive action current is shown on the right for each pair (green trace: EGFP(+) cell; black trace: EGFP(-) cell). **C and D**) summary plots of the results (n=18 pairs). **C**) average number of spikes evoked within the 200 ms following the stimulation. **D**) amplitudes of the average capacitive currents evoked by the stimulation. **E)** c-fos expression in activated glomeruli in vivo. **E**) c-Fos expression in PG interneuron subtypes (CB+, CR+, TH+) in activated glomeruli *in vivo*. Inserts show higher magnifications of select region (boxes on overviews). The graph shows the proportion of each PG interneuron subtypes expressing c-Fos. Scale bars: 25 µm.

Finally, we also investigated the activity of CR(+) PG cells *in vivo* using the expression of the immediate early gene c-Fos as a marker of cellular activation (**Fig.6D**). We compared the relative activation of various PG cell subtypes under basal conditions in mice housed in their regular cage. In these animals, occasional glomeruli were heavily labeled by a c-Fos immunostaining (n=1104 c-Fos+ cells in 3 mice). In these glomeruli, the majority of CB(+) PG interneurons were positive for c-Fos (72.3 ± 2%), in line with their pronounced activity *in vitro* and *in vivo* in anesthetized animals (Najac et al., 2015). In comparison, a smaller percentage of TH(+) PG interneurons were positive for this immediate early gene (28.1 ± 7%), while only rare CR(+) c-Fos(+) PG cells were observed (11.5 ± 1%) consistent with their low network-driven activity in slices. Thus, all together, our data suggest that the physiological properties of CR(+) PG cells limit their contribution in olfactory bulb circuit processing.

### DISCUSSION

Our study shows that CR+ PG cells, independent of their spatial origin and age, receive little synaptic inputs and do not fire or fire at most a single and often small action potential in response to stimuli that strongly activate other PG cell subtypes. Altogether, the synaptic and intrinsic properties of CR+ PG cells resemble those of immature neurons not yet integrated in the pre-existing network. Surprisingly, however, our birth dating experiments indicate that CR+ PG cells remain in this immature stage for weeks, if not months, as if they would never fully mature. Thus, CR+ PG cells constitute a particularly large pool of latent neurons that unlikely participate in olfactory bulb circuit computations.

### Calretinin defines a homogeneous population of PG interneurons with complex spatio-temporal origins

Our long term fate mapping of neural stem cells located in defined SVZ walls confirms the dual origin of CR(+) PG interneurons from the dorsal and the septal SVZ throughout postnatal life (Fernández et al. 2011) and extent it to adulthood. The persistent dorsal origin of CR(+) PG interneurons observed at P90 is consistent with a previous fate mapping study reporting that a fraction of these cells derives from the pallium (Emx1 lineage), which gives rise to the dorsal part of the postnatal SVZ (Kohwi et al. 2007). This previous report also showed that a fraction of CR(+) PG interneurons derives from the Dlx5/6 lineage, a transcription factor that is strongly active in the embryonic lateral ganglionic eminence and in the septum which give rise to the lateral and septal parts of the postnatal SVZ, respectively. Our observations indicate that CR(+) PG interneurons from the Dlx5/6 lineage originate from the septal most regions of the postnatal SVZ since they were consistently absent following lateral SVZ EPO at both postnatal and adult stages. The rapid decrease of germinal activity observed in the septal wall during the first 15 days of postnatal life suggests, however, that the proportion of CR(+) interneurons originating from the dorsal SVZ gradually increases over time, consistent with the gradual increase in the proportion of CR(+) interneurons originating from the Emx1 lineage with age (Kohwi et al. 2007).

We demonstrate that this complex spatial and temporal origin does not translate into morphological and electrophysiological differences. Using the CR::EGFP reporter mouse line, we found that CR(+) PG cells originating from distinct stem cells and generated at different developmental ages have rather uniform properties. The variability of some of these properties likely reflects different stages of maturation rather than different origins. Yet, EGFP labels only half of the CR-expressing PG cells in adult mice, leaving open the possibility that unlabeled CR(+) PG cells form a different subpopulation of neurons with more mature properties and a possibly different impact on the glomerular network. Several observations, however, suggest that the coexistence of two functionally distinct CR(+) PG cell populations is unlikely. First, we demonstrate that EGFP is expressed for at least two months in CR(+) PG cells, unlike in other transgenic models in which newborn neurons transiently express a reporter during a defined period of their maturation (Overstreet et al., 2004; Spampanato et al., 2012). Second, we show that the morphology of EGFP(+) and EGFP(-) CR(+) PG cells is largely similar. Third, PG cells that do not fire in response to OSN stimulation represent around 40% of the PG cells in our previous blind recordings, consistent with the abundance of CR(+) PG cells relative to other subtypes (Najac et al., 2015). Fourth, a recent study reported similar intrinsic membrane properties in CR(+) PG cells expressing EGFP in another transgenic strain (Fogli Iseppe et al., 2016). Thus, despite their incomplete labeling in the transgenic mouse used in our study, CR(+) PG cells likely constitute a population of neurons with homogeneous physiological properties.

### CR(+) PG cells show properties of immature neurons that persist over time

Several subtypes of neurons continue to be produced after birth. In the olfactory bulb, a majority of inhibitory granule cells are produced in parallel to a smaller population of various PG interneurons. Neurogenesis also persists in the hippocampus from stem cells located in the subgranular zone of the dentate gyrus that produce glutamatergic granule cells. The maturation and integration of these newborn neurons in the pre-existing network has been examined in great details in both structures and, despite small timing differences and specific synaptic connections, follows essentially the same sequence of morphological and functional maturation (reviewed in (Lepousez et al., 2015; Toni and Schinder, 2015)). Interestingly, the properties of CR(+) PG cells described in this paper are reminiscent of those of immature newborn neurons not yet fully integrated in the preexisting olfactory bulb or hippocampal network, i.e. 2 to 4 weeks after their birth (Carleton et al., 2003; Esposito et al., 2005). Like them, they have a large electrical input resistance and express a low density of voltage-activated channels. In addition, they fire at most a single action potential and many cells are not capable of firing a full size action potential, a property observed in newborn neurons up to 3 weeks after their birth. They also receive little synaptic inputs compared to resident interneurons embedded in the same network. Finally, their excitatory synapses are enriched with NMDA receptors, a characteristic of immature and plastic excitatory synapses during postnatal development or during maturation in adult-generated newborn neurons (Ge et al., 2007; Grubb et al., 2008; Katagiri et al., 2011). However, these immature properties persist for extended periods of time in CR(+) PG interneurons, greatly contrasting with the rapid activity-dependent maturation and integration (i.e. 4-6 weeks) of other populations of newborn bulbar or hippocampal interneurons (Belluzzi et al., 2003; Carleton et al., 2003; Esposito et al., 2005; Livneh et al., 2014).

### CR+ PG cells unlikely participate to circuit function

Type 2 PG cells are activated by mitral and tufted cells and in turn collectively produce a strong feedback inhibition of these principal neurons (Najac et al., 2011; Shao et al., 2012; Shao et al., 2013; Najac et al., 2015; Geramita and Urban, 2017). Thus, OSN stimulation produces a barrage of unsynchronized IPSCs in mitral and tufted cells whose time course matches well with the burst of action potentials fired in other type 2 PG cells (Najac et al., 2015; Geramita and Urban, 2017). Current models suggest that this intraglomerular inhibition serves as a gatekeeper that shunt weak OSN inputs (Gire and Schoppa, 2009) and differently modulates mitral and tufted cells, thereby separating their firing phase along the respiration cycle (Fukunaga et al., 2012; Fukunaga et al., 2014) and generating distinct modes of intensity coding in these two output channels (Geramita and Urban, 2017). These afferent-driven functions require functional excitatory input and inhibitory output connections as well as the ability to fire repetitively and are therefore not compatible with the properties of CR(+) PG cells. Even the small proportion of CR(+) PG interneurons that receive a significant amount of excitatory synaptic inputs, and which likely represents their most advanced stage of maturation, cannot fire more than once in response to the temporally complex and prolonged depolarization induced by mitral and tufted cells (Fogli Iseppe et al., 2016). Activation of these most “mature” CR(+) PG cells should therefore produce, at most, a single IPSC in their postsynaptic target. However, the synaptic targets of CR(+) PG cells are still unknown and a phasic OSN-driven inhibition that cannot adapt to the circuit’s afferent activity has, to our best knowledge, not yet been observed in the glomerular network.

The capacity of CR(+) PG cells to release GABA at output synapses has also to be demonstrated. Self-inhibition is a read-out for GABA release in many PG cells (Smith and Jahr, 2002; Murphy et al., 2005; Maher and Westbrook, 2008) but CR(+) PG cells do not generate self-inhibition after step depolarization (our data, not shown). Moreover, ultrastructural data supporting the existence of functional output synapses from CR(+) PG cells are also sparse in the literature compared to those concerning CB(+) and TH(+) cells. Previous works indicate that CR(+) PG cells, quote, *appear to establish fewer synapses than CB(+) and TH(+) cells* (Panzanelli et al., 2007). Moreover, a surprisingly large fraction of CR(+) PG cells is immuno-negative for GAD/GABA (Kosaka and Kosaka, 2007) or VGAT, the vesicular transporter for GABA (Sawada et al., 2011) whereas 100% of CB(+) and TH(+) subtypes express these markers. Thus, the low connectivity, the stereotyped firing pattern and the uncertain GABA output of CR(+) PG cells suggest that their contribution to olfactory bulb processing is, at most, limited.

Altogether, our results reveal the atypical and intriguing properties of CR(+) PG interneurons. The function of this large neuronal population with complex spatial and temporal origin and that is continuously produced remains, however, elusive. We have not examined the possibility that CR(+) PG cells are activated by other, extraglomerular pathways and our results do not exclude the possibility that the small fraction of CR(+) PG cells that fire and receive significant inputs from mitral and tufted cells inhibit specific targets within the glomerular network. It is tempting to speculate that the abundant population of immature CR(+) cells constitutes a reserve pool of latent and not fully differentiated interneurons that could supply the glomerular network on demand, based on specific sensory experience. Such recruitment, that may take the form of a functional maturation or a fate conversion in other PG cell subtypes, would provide an unsuspected extra level of plasticity to the glomerular network and would compensate the lower postnatal neurogenesis of PG cells compared to granule cells.

## METHODS

### Animals

The Rosa^YFP^ mice express YFP upon Cre recombination (Srinivas et al., 2001) (line 006148; Jackson laboratory). The CR::EGFP mice express EGFP under the control of the CR promoter (Caputi et al., 2009). The Kv3.1::EYFP mice express EYFP under the control of the Kv3.1 K^+^ channel promoter (Metzger et al., 2002).

### Acute slice preparation

All experimental procedures conformed to the French Ministry and local ethics committee (CREMEAS) guidelines on animal experimentation. Transgenic mice of different ages were decapitated and the olfactory bulbs removed and immersed in ice-cold solution containing (in mM) 83 NaCl, 26.2 NaHCO_3_, 1 NaH_2_PO_4_, 2.5 KCl, 3.3 MgSO_4_, 0.5 CaCl_2_, 70 sucrose, and 22 D-glucose, pH 7.3 (300 mOsm/l), bubbled with 95% O_2_/5% CO_2_. Horizontal olfactory bulb slices (300 µm) were cut using a Microm HM 650V vibratome (Microm, Walldorf, Germany) in the same solution. The slices were incubated for 30 min at 34˚C and then stored until use at room temperature in artificial cerebrospinal fluid (ACSF) containing (in mM) 125 NaCl, 25 NaHCO_3_, 2.5 KCl, 1.25 NaH_2_PO_4_, 1 MgCl_2_, 2 CaCl_2_, and 25 D-glucose, continuously oxygenated with 95% O_2_/5% CO_2_.

### Electrophysiology

Experiments were conducted at 32-34°C under an upright microscope (SliceScope, Scientifica, Uckfield, UK) with differential interference contrast (DIC) optics. Loose cell-attached recordings (15-100 MΩ seal resistance) were made with pipettes filled with ACSF. OSN axons bundles projecting inside a given glomerulus were stimulated using a theta pipette filled with ACSF as previously described (Najac et al., 2011). The electrical stimulus (100 µs) was delivered using a Digitimer DS3 (Digitimer, Welwyn Garden City, UK). Most whole-cell recordings were made with pipettes (3-6 MOhm) filled with (in mM): 135 K-gluconate, 2 MgCl2, 0.025 CaCl2, 1 EGTA, 4 Na-ATP, 0.5 Na-GTP, and 10 HEPES, pH 7.3 (280 Osm/L). Atto 594 fluorescent dye (5 mM, Sigma Aldrich, St. Louis, MO) was routinely added to this internal solution to confirm the glomerular projection of the recorded cell. Voltages indicated in the paper were corrected for the junction potential (−15 mV). Sodium currents were recorded using a Cs-based internal solution, containing (in mM): 120 Cs-MeSO3, 20 tetraethylammonium-Cl (TEA), 5 4-aminopyridine (4-AP), 2 MgCl2, 0.025 CaCl2, 1 EGTA, 4 Na-ATP, 0.5 Na-GTP, and 10 HEPES, pH 7.3 (280 Osm/L, 10 mV junction potential), and in the presence of CdCl (100 µM), TEA (20 mM) and 4-AP (5 mM) in the bath. The tip of the recording pipette was systematically coated with wax and we used the capacitance neutralization of the amplifier in order to reduce the capacitance of the pipette. In current-clamp recordings, a constant hyperpolarizing current was injected in order to maintain the cell at a potential of −60/-70 mV. Without any current injected, the membrane potential remained at positive potentials. In voltage-clamp recordings, the access resistance was compensated by 50 to 80%.

Recordings were acquired with a multiclamp 700B amplifier (Molecular Devices, San Jose, CA), low-passed filtered at 2-4 kHz and digitized at 10 kHz using the AxoGraph X software (Axograph Scientific).

### Recording analysis

Data were analyzed using Axograph X (Axograph Scientific). Current amplitudes were measured as the peak of an average response computed from multiple sweeps. Spike amplitudes were measured between the peak and the most negative voltage reached immediately after the spike. Membrane time constants were obtained from the exponential fit of the voltage deflection caused by a small hyperpolarizing current. Individual spontaneous excitatory (EPSC) or inhibitory (IPSC) currents with amplitude >5 pA were automatically detected by the Axograph X software using a sliding template function. For measuring the AMPA/NMDA ratio, neurons were first voltage-clamped at –75 mV and their excitatory inputs were evoked with an electrical stimulation in the EPL. Stimulation intensity was adjusted to obtain isolated responses. Neurons were then voltage-clamped at +50/60mV to release the Mg^2+^ blockade of NMDA receptors. After a stabilization period of 5 to 15 min, the NMDA receptor antagonist D-2-amino-5-phosphonopentanoic acid (D-AP5, 50 μM, Abcam biochemicals, Cambridge, UK) was bath applied to isolate AMPA currents. Isolated AMPA currents were digitally subtracted offline from the total EPSCs to obtain the NMDA currents.

### Plasmids

Plasmids pUb-TdTomato plasmid (gift from Prof D. Jabaudon) and pCAG-CRE (#13775, Addgene, Cambridge, MA, USA) were prepared with a PEG-LiCl protocol and used at a concentration of 5 µg/µL. For EPO, plasmids were mixed 1:10 with a contrast solution of fast green (0.2%) in sterile PBS.

### Electroporations

Electroporations (EPO) were performed in postnatal 2 days (P2) pups as described previously (Boutin et al., 2008; Fernandez et al., 2011). Briefly, pups were anesthetized in ice and placed on a custom-made support in a stereotaxic rig. Injections were performed at the midpoint of a virtual line traced between the eye and the lambda. A 34G needle attached to a Hamilton syringe was inserted at a depth of 2.5 mm from the skull surface and 1.5µl of plasmid solution was injected into the lateral ventricle. The accuracy of the injection was verified by the filling of the ventricle with the contrast solution. Successfully injected mice were then subjected to 5 electrical pulses (95 V, 50 ms, separated by 950 ms intervals) using the Super Electroporator NEPA21 type II (Nepa Gene Co., Ltd, Ichikawa-City, Japan) and tweezer electrodes coated with conductive gel (SignaGel, Parker Laboratories, Fairfield, NJ, USA). Electrodes were positioned in order to target the lateral, the dorsal or the septal wall. After EPO, pups were warmed up until they fully recovered and returned to their mother.

### 5-bromo-2-deoxyuridine (BrdU) treatments and tissue processing

For analysis of CR(+) PG cells spatial and temporal origin, BrdU (Sigma-Aldrich B5002) was added in drinking water at a concentration of 1 mg/mL supplemented with sucrose at 10 mg/mL for 15 days (from P12 to P27), protected from light and changed every 2 days. Animals were sacrificed at 45 days post EPO. After deep anesthesia with intraperitoneal pentobarbital, mice were transcardially perfused with a Ringer solution and a 4% paraformaldehyde in 0.1M PB solution, brains were removed, post-fixed for 24h and cut with a vibratome (Leica VT1000S, Wetzlar, Germany). 50 µm coronal sections were collected from the olfactory bulb to the lateral ventricle in series of 6 and kept at −20° C in antifreeze solution (glucose 15%, sodium azide 0,02%, ethylene glycol 30%, PB 0,1M).

For birth dating of CR(+) PG cells, CR::EGFP transgenic mice (P30) received four intraperitoneal injections of 20 mg/mL BrdU in sterile 0.9% NaCl solution (100 mg/kg), 2 hours apart. At days 15, 20, 30, 40, 50 and 60 after injection, mice were anesthetized by intraperitoneal injection of a ketamine and xylazine mixture (90 and 100 mg/kg, respectively), and transcardially perfused with 4% paraformaldehyde diluted in PBS. The olfactory bulbs were removed and postfixed overnight at 4 ˚C. Horizontal sections (30 µm) were sliced on a vibratome (Leica VT1000S).

### Immunostaining

For BrdU staining, free-floating sections were incubated for 1 hr in TritonX100 (0.2-0.4% in PBS), pretreated in 1N (10 min, 37˚C) and 2N (20 min, 37˚C) HCl, and washed in 0.1 M borate buffer pH 8.4 (5-30 min). After rinsing three times with PBS (10 min), sections were incubated in blocking solution containing BSA (2%) and TritonX100 (0.3%), for 1 hr before incubation with the primary antibodies (see table 1).

**Table 1:**

| Primary antibodies | Species | Concentration | Source (catalog number) |
| --- | --- | --- | --- |
| Anti-BrdU | Rat | 1:500 | ABD Serotech (ABD0030) |
| Anti-BrdU | Rat | 1:500 | Abcam Ab6326 |
| Anti-Calretinin | Rabbit | 1:1000 to 1:2000 | SWANT (7697) |
| Anti-Calretinin | Mouse | 1:2000 | SWANT (6B3) |
| Anti-Calbindin | Mouse | 1:2000 | SWANT (D28K) |
| Anti-cFos | Goat | 1:50 | Santa Cruz (Sc-52-G) |
| Anti-TH | Mouse | 1:400 | Millipore (MAB318) |
| Anti-GFP | Chicken | 1:1000 | JAVES (GFP-1020) |
| Anti-Doublecortin | Goat | 1:500 | Santa Cruz (Sc-8066) |
| Anti-RFP | Rabbit | 1:1500 | MBL (PM005) |
| **Secondary Antibodies** | **Specie** | **Concentration** | **Source (catalog number)** |
| Anti-rat AlexaFluor 555 | Donkey | 1:1000 | ThermoFisher (A-21434) |
| Anti-rat AlexaFluor 546 | Goat | 1:200 | ThermoFisher (A-11081) |
| Anti-mouse AlexaFluor 555 | Donkey | 1:1000 | ThermoFisher (A-31570) |
| Anti-mouse AlexaFluor 647 | Donkey | 1:1000 | ThermoFisher (A-31571) |
| Anti-rabbit AlexaFluor 555 | Donkey | 1:1000 | ThermoFisher (A-31572) |
| Anti-rabbit AlexaFluor 647 | Donkey | 1:1000 | ThermoFisher (A-31573) |
| Anti-rabbit AlexaFluor 633 | Goat | 1:200 | ThermoFisher (A-21070) |
| Anti-goat AlexaFluor 555 | Donkey | 1:1000 | ThermoFisher (A-21432) |
| Anti-chicken biotinylated | Donkey | 1:1000 | Jackson Lab (703-065-155) |
| Streptavidin-DTAF | - | 1:250 | Jackson Lab (016-010-084) |

For cFos staining, antigen retrieval treatment was performed in 10 mM Citrate Buffer pH 6 (20 min, 80°C). After rinsing three times with TritonX100 (0.4% in PBS), sections were incubated in blocking solution for 2-4 hours at room temperature (BSA 0.25%, Casein, 0.05%, Top block 0.25% + TritonX100 0.4% in PBS) before incubation with the primary antibodies overnight at 4°C (table 1).

Following incubation with primary antibodies, sections were rinsed then incubated 2h at room temperature with the corresponding secondary antibody (table 1). To amplify YFP signal, sections were incubated with a biotinylated secondary antibody followed by streptavidin-DTAF for 45min. For morphological reconstruction, the tomato signal was amplified with an anti-RFP antibody. Finally, sections were counterstained with DAPI and mounted with Prolong Antifade (Thermosfisher, Waltham, MA, USA) or Fluoromount (Southernbiotech, Birmingham, AL, USA) mounting medium.

### Quantifications

For the analysis of CR(+) PG interneurons neurogenesis and spatial and temporal origin, YFP(+) cells were counted in the glomerular layer on live images using an epifluorescent microscope (Leica DM5500, objective HCX PL APO 40x 1,25 oil).

For CR::EGFP interneurons origin, 3 animals were analyzed for dorsal and septal EPO, and the number of CR(+)and EGFP(+) PG cells among the Td-Tomato-expressing population analyzed (>85 Td-tomato(+) cells per EPO).

For CR(+)/EGFP(+) PG interneurons birth dating analysis, series of 3 slices (3 pictures per slice) were analyzed with a confocal microscope (Leica SP5), and the number of BrdU-positive nuclei and EGFP-positive cells located in the glomerular layer counted (>120 BrdU-positive cells per animal).

For cFos analysis, images were taken with a confocal microscope (Leica SP5) using a 40x objective (HCX PL APO 40x 1.25 oil). cFos-positive nuclei expressing CR, CB and TH in the glomerular layer were quantified (>120 cFos-positive cell per animal; 3 animals per condition). Analysis was focused on cells bordering glomeruli where an intense cFos staining was observed.

For arborization analysis of CR(+) PG interneurons, 0.3 µm stack images were taken with a confocal microscope (Leica SPE), and 3D neurons reconstruction were performed and analyzed with the software Neurolucida 360 (43 EGFP(+) cells vs 12 EGFP(-), equally distributed between septal and dorsal EPO, i.e. 26 and 29 CR(+) neurons, respectively; 3 mice per EPO). For a direct comparison with CB(+) PG interneurons, CB(+) neurons were reconstructed from 3 dorsally electroporated mice, as they were consistently absent from septally electroporated ones.

Results are expressed as mean ± SEM.

Statistical significance was determined by unpaired Student’s *t* test at the *p*< 0.05 level or a Mann-Whitney test when data did not assume a normal distribution.

## Acknowledgements

Work in the DD lab was supported by the Centre National pour la Recherche Scientifique, the Université de Strasbourg and the Agence Nationale pour la Recherche (Grant ANR-12-JSV4-006-01). ASD was funded by a fellowship from the Ministère de la Recherche and by a fellowship from the Fonds Paul Mandel pour les Recherches en Neuroscience. Work in the OR lab was supported by a grant from the “Programme Avenir Lyon Saint-Etienne”. We thank Sophie Reibel-Foisset and the animal facility Chronobiotron (UMS 3415 Centre National pour la Recherche Scientifique and Université de Strasbourg), the Plateforme Imagerie In Vitro-Neuropôle-Strasbourg, Bastien Leclerc and Aline Huber for their technical assistance. We also thank Linda Overstreet-Wadiche, Cécile Viollet and members of the lab for their advice and support.

**Authors contributions**
NB, EG, ASD, LF, SK and DDSJ did the experiments. DDSJ and OR designed the project and wrote the paper that was edited by all the authors

